# Single-cell Transcriptomic Profiling of Pancreatic Ductal Adenocarcinoma: Epithelial Reprogramming and Systemic Immune Exhaustion

**DOI:** 10.1101/2025.11.10.687730

**Authors:** Qinyu Xu

**Affiliations:** Zu Chongzhi Center, Duke Kunshan University, 8 Duke Avenue, Kunshan, 215316, Jiangsu, China

**Keywords:** Pancreatic ductal adenocarcinoma, Single-cell RNA sequencing, Epithelial reprogramming, Immune exhaustion, Tumor microenvironment

## Abstract

Pancreatic ductal adenocarcinoma (PDAC) remains one of the most lethal human malignancies, with a five-year survival rate below 10%, underscoring the urgent need to unravel its complex pathophysiology. In this study, we employ an integrative single-cell RNA sequencing (scRNA-seq) approach to comprehensively characterize both the pancreatic tumor microenvironment and peripheral blood mononuclear cells (PBMCs) from PDAC patients and healthy controls. Using Harmony for batch correction and Azimuth for automated cell-type annotation, we identify cell-type–specific transcriptional alterations through differential expression and functional enrichment analyses. Our results reveal a system-wide disruption involving both epithelial and immune compartments. Epithelial cells exhibit a “de-skilling” phenotype, losing digestive enzyme expression while activating metabolic and ribosomal programs that sustain malignant proliferation. The local immune microenvironment displays chronic inflammation and remodeling, whereas systemic immune exhaustion is evident in cytotoxic T, NK, and myeloid populations, marked by impaired effector activity and metabolic imbalance. Collectively, these findings depict PDAC as a coordinated multi-compartmental failure encompassing epithelial reprogramming, local immune dysregulation, and systemic immune collapse, providing a holistic single-cell view of PDAC pathogenesis and offering mechanistic insights that may guide biomarker discovery and the development of targeted therapeutic strategies.

## 1 Introduction

Pancreatic cancer is recognized as one of the most lethal malignancies with an increasing mortality rate worldwide, reflecting its exceptionally high fatality [1]. The global five-year survival rate remains below 10%, primarily due to late-stage diagnoses and the lack of an effective screening method [2, 3]. The increasing incidence and mortality are further exacerbated by aging populations, obesity, diabetes, and disparities in healthcare access, highlighting the importance of intensified research into screening strategies, molecular mechanisms, and novel therapeutics [1, 4]. Collectively, the aggressive biology, late presentation, and poor survival outcomes of pancreatic cancer establish it as a critical public health priority requiring sustained global attention.

Given the extremely high lethality and poor prognosis of cancers such as pancreatic cancer, understanding the role of immune cells within the tumor microenvironment has become increasingly important. Immune cells are central players in cancer biology, capable of both suppressing and promoting tumor progression depending on their subtype, localization, and functional state. For instance, cytotoxic CD8^+^ T cells and Th1 CD4^+^ T cells can recognize and eliminate malignant cells, and their infiltration is often associated with favorable patient outcomes [5, 6]. Dendritic cells, particularly cDC1, link innate and adaptive immunity by presenting tumor antigens and initiating T cell responses, thereby enhancing the effectiveness of immunotherapy [7, 8]. Importantly, the prognostic impact of immune infiltration is cancer-type specific. For instance, while in melanoma or lung cancer higher immune infiltration generally predicts longer survival, in gliomas or uveal melanoma it may portend worse outcomes [9]. Collectively, these findings underscore that immune cells are not only biomarkers of prognosis but also critical targets for next-generation cancer therapies, making their study indispensable in the broader effort to improve outcomes for lethal cancers such as pancreatic cancer.

Building upon the urgent need to address lethal cancers and the complex roles of immune cells in shaping tumor progression, single-cell sequencing technologies have emerged as essential tools in cancer research. Single-cell RNA sequencing (scRNA-seq) enables the dissection of individual tumor and immune cell states, providing unprecedented resolution of the tumor microenvironment [10]. This is particularly crucial in cancers like pancreatic cancer, where the dense stroma, immunosuppressive milieu, and heterogeneous cell populations contribute to therapeutic resistance and poor prognosis [11]. By profiling immune subsets, researchers can map immune landscapes that correlate with patient outcomes and identify novel therapeutic targets [12]. As cancer research moves toward personalized interventions, single-cell sequencing stands at the forefront, enabling a deeper mechanistic understanding of both tumor biology and immune dynamics [13, 14]. Ultimately, its integration into oncology research offers a path to overcome the limitations of current therapies and to improve outcomes for patients with aggressive cancers such as pancreatic cancer [11].

In this study, we perform an integrated single-cell RNA sequencing analysis for both pancreatic tissue and peripheral blood mononuclear cell (PBMC) samples from patients with pancreatic ductal adenocarcinoma (PDAC) and healthy controls. Starting with rigorous quality control, low-quality cells are effectively removed based on mitochondrial gene proportion and the number of detected features. Accordingly, for each sample type, we apply Harmony integration to correct for batch effects and enable robust cross-sample comparison. Cell types are systematically annotated using the Azimuth reference-based pipeline, which allows for precise characterization of cellular compositions. We then conduct differential gene expression analysis between cancer and healthy groups within specific cell populations, visualized through volcano plots, and follow this with functional enrichment analysis to uncover dysregulated biological pathways. In addition, we construct cell trajectories for some PBMC cell types, enabling us to explore where cancerous and normal cells are positioned along the pseudotime axis of cellular progression and to identify potential subtypes associated with distinct developmental states.

This multi-faceted approach provides a comprehensive view of the transcriptional alterations in both the pancreatic tumor microenvironment and systemic immune circulation, offering valuable insights into the molecular mechanisms of PDAC pathogenesis and potential biomarkers for early detection or therapeutic targeting.

## 2 Methods

### 2.1 Quality Control of Single-Cell RNA Sequencing Data

The raw single-cell RNA sequencing data are obtained from the Gene Expression Omnibus (GEO) database under accession number GSE155698. Prior to data integration, rigorous quality control (QC) is performed on each individual sample to ensure data reliability and remove technical artifacts. The QC pipeline is implemented using the Seurat package in R.

For each sample, raw gene expression matrices are initially loaded and converted into Seurat objects, and key QC metrics are calculated, including the number of unique genes detected per cell (`nFeature_RNA`), the total number of molecules detected per cell (`nCount_RNA`), and the percentage of mitochondrial gene expression (`percent.mt`). Cells are then filtered based on pre-defined thresholds to remove low-quality cells and potential doublets: specifically, cells with a mitochondrial gene percentage ≥ 5% are excluded to remove damaged or dying cells, and cells with fewer than 200 or more than 2500 detected genes are removed to eliminate empty droplets and multiplets.

This stringent QC process ensures that only high-quality cells are retained for subsequent analyses, thereby reducing technical noise and improving the robustness of downstream integration and differential expression results. Following QC, filtered count matrices are regenerated in standard 10X or h5 format for consistent processing across all samples.

### 2.2 Data Integration

The integration of single-cell RNA sequencing data from multiple samples after quality control is a critical step to address technical batch effects, which can obscure biological signals and confound downstream comparative analyses. To achieve a robust and accurate integration of our dataset, we employ the Harmony algorithm, which is recognized for its efficiency in removing dataset-specific biases while preserving true biological heterogeneity [15]. The entire process is implemented using the Seurat package (v5) in R [16], following a structured pipeline outlined below.

#### Data Merging and Initial Object Creation

The analysis begins with the creation of individual Seurat objects for each sample. The input data for each sample consists of three core components: a vector of cell barcodes, a vector of all detected genes, and a gene expression matrix where entries represent the count of mRNA molecules for each gene (row) in each cell (column). These components are integrated into a standardized data structure using the `CreateSeuratObject` function, which encapsulates the raw count matrix along with cell-level and feature-level metadata. Subsequently, all individual Seurat objects are merged into a single unified object using the `merge` function. This initial merging step is fundamental, as it consolidates data from all sources into a single entity for coordinated processing, while preserving the sample origin of each cell in the `orig.ident` metadata field for later batch correction.

#### Pre-processing and Dimensionality Reduction

Prior to integration, the merged dataset undergoes standard pre-processing procedures. This involves normalizing the gene expression data using the `NormalizeData` function to mitigate technical artifacts and sequencing depth biases, thereby ensuring comparability across samples. Subsequently, highly variable features are identified via `FindVariableFeatures` to capture genes that drive biological heterogeneity. Furthermore, we scale the data using `ScaleData` to regress out sources of variation like sequencing depth. Linear dimensionality reduction is then performed via Principal Component Analysis (PCA) using the `RunPCA` function. The top 30 principal components capture the major sources of variation in the data, serving as a condensed representation for the subsequent integration step.

#### Harmony Integration for Batch Correction

The core integration is performed using Harmony via the `IntegrateLayers` function with `method = HarmonyIntegration`. This function takes the PCA embedding as input and iteratively corrects it. The algorithm uses a soft k-means clustering approach to group cells by their cell type (biological signal) while simultaneously applying a correction factor to minimize the influence of batch origin (technical signal). The output is a new, batch-corrected low-dimensional embedding named `“harmony”`. The significance of this step is that it enables cells of the same biological type from different samples to co-localize in the embedding space, which is a prerequisite for accurate cross-sample comparisons.

#### Downstream Clustering and Visualization

Following integration, all down-stream analyses are performed on the corrected Harmony embeddings. Cell-to-cell relationships are quantified by first constructing a shared nearest neighbor (SNN) graph using the `FindNeighbors` function, which computes the k-nearest neighbors for each cell based on the Harmony-corrected space. The `FindClusters` function then applies the Louvain algorithm to this graph to identify distinct cell populations. Finally, to visualize the high-dimensional integrated data in two dimensions, we perform non-linear dimensionality reduction using Uniform Manifold Approximation and Projection (UMAP) with the `RunUMAP` function. UMAP works by modeling the local relationships between cells in the Harmony space and finding a low-dimensional layout that preserves this topological structure. The resulting plot provides an intuitive representation of the global cell landscape, where the proximity of cells reflects their transcriptional similarity after the removal of batch effects.

### 2.3 Cell Type Annotation

Following data integration, automated cell type annotation is performed using Azimuth, a web-based tool that leverages a pre-defined, curated reference atlas to transfer labels to the cells of a query dataset [17]. This reference-based approach is superior to unsupervised clustering, as it ensures annotation consistency, leverages existing biological knowledge, and allows for the identification of rare or transient cell states that might be overlooked otherwise.

#### Reference-Based Annotation with Azimuth

Azimuth’s annotation engine is built upon the Seurat framework and utilizes a robust label transfer methodology [18]. The core mechanism is a two-step process. Firstly, the algorithm finds “anchors” between the query and reference datasets and constructs an SNN graph. A query cell’s label is predicted by performing a weighted vote among its closest reference neighbors in this graph, where the weight is a function of the SNN graph edge strength. This method is more robust to technical noise than simple distance-based approaches [17]. Secondly, for each cell and each possible cell type, Azimuth calculates a prediction score that reflects the confidence of the assignment. This score is derived from the weights of the edges connecting the query cell to its reference neighbors of that specific type. A score close to 1 indicates that the query cell is strongly and exclusively connected to reference cells of the assigned type. Additionally, the mapping score is computed, which measures the aggregate strength of a cell’s connections to *all* reference cells. A low mapping score suggests the cell is a poor match to the entire reference atlas, potentially indicating a low-quality cell, doublet, or a novel cell type not present in the reference.

#### Data Preparation for Azimuth Compatibility

A critical data preparation step is required to bridge the compatibility gap between our analysis environment and the Azimuth web service. Our integrated dataset is processed using Seurat v5, which employs a modern `Assay5` object utilizing a unified `layers` architecture for efficient data storage. In contrast, the Azimuth server is built upon Seurat v4, which relies on a legacy data structure where counts and normalized data reside in distinct slots of a standard `Assay` object. To resolve this version mismatch, we implement a custom conversion function. This function programmatically detects the `Assay5` object, consolidates the data from its disparate layers into unified count and normalized data matrices using `JoinLayers`, and then reconstructs a new, v4-compatible `Assay` object. Subsequently, the `DietSeurat` function is applied to streamline the object by removing all non-essential components, retaining only the `RNA` assay and the `pca` dimensional reduction. This process yields a significantly smaller file that is fully compatible with Azimuth’s data input requirements, enabling robust and efficient automated cell type prediction.

#### Quality Filtering of Annotations

Azimuth returns a comprehensive annotation table for each cell. The PBMC samples are annotated with a hierarchical classification system (Level 1, Level 2, and Level 3), providing resolution from major lineages (e.g., T cells, Myeloid cells) to fine subsets (e.g., CD4 Naive, CD8 TEM). The pancreatic tissue samples are annotated with a single-level classification. To ensure the robustness and reliability of downstream analyses, we perform a stringent quality filter on the annotated cells. This step removes cells with low-confidence annotations, which could introduce noise and spurious results.

For PBMC samples, a cell is retained only if its prediction scores for all three hierarchical levels (`predicted.celltype.l1`, `predicted.celltype.l2`, `predicted.celltype.l3`) are greater than 0.7 and its `mapping.score` is greater than 0.5. For pancreatic tissue samples, a cell is retained only if its `predicted.annotation.l1` score is greater than 0.7 and its `mapping.score` is greater than 0.5.

Cells that do not meet these criteria are filtered out. This rigorous filtering strategy ensures that only cells with high-confidence annotations are carried forward, thereby increasing the credibility and interpretability of subsequent differential gene expression analyses. The final, filtered dataset is used for all further investigations.

### 2.4 Differential Expression and Functional Enrichment Analysis

To identify molecular signatures specific to PDAC within defined cell populations, we perform differential expression (DE) analysis followed by functional enrichment analysis. This multi-step approach allows for the identification of not only dysregulated genes but also the biological processes they implicate.

#### Cell-Type-Specific Differential Expression Analysis

Differential expression analysis is conducted to compare gene expression profiles between cancer (PDAC) and healthy control samples within each cell type. For a given cell type of interest, we first create a new metadata column, `celltype.condition`, by concatenating the annotated cell type and the condition (i.e., `“Cancer”` and `“Healthy”`). The cell identities in the Seurat object are then set to this new column using `Idents()`. Differential expression testing between the two conditions is carried out using the `FindMarkers` function with default parameters, which employs a Wilcoxon rank sum test.

#### Visualization with Volcano Plots

The results of the differential expression analysis were visualized using volcano plots, a standard visualization that simultaneously displays the statistical confidence and biological effect size for thousands of genes. In this plot, the x-axis represents the log2 fold change (log2FC), which quantifies the magnitude of expression difference between the two conditions (i.e., `“Cancer”` VS `“Healthy”`). A log2FC of 0 indicates no change, a positive value indicates higher expression in the first condition (i.e., `“Cancer”`), and a negative value indicates lower expression. The y-axis represents the −log10 of the adjusted p-value, which is a transformation of the statistical significance that accounts for multiple testing. A higher value on the y-axis indicates greater statistical significance. This transformation enlarges the visual distance between highly significant and non-significant genes, making it easier to identify key candidates.

The plot is generated using the `ggplot2` package. Genes with an average log2 fold change (avg log2FC) greater than 0.6 or less than −0.6 and an adjusted p-value (p val adj) less than 0.05 are considered as significantly up-regulated or down-regulated, respectively. These thresholds are represented by vertical dashed lines at *x* = ±0.6 and a horizontal dashed line at *y* = −*log*_10_(0.05) ≈ 1.3. Significantly up-regulated genes are colored in red, down-regulated genes in cyan, and non-significant genes in grey. To highlight the most biologically relevant findings, the top 5 most significant (lowest adjusted p-value) genes from both the up-regulated and down-regulated gene sets are automatically labeled on the plot using the `geom_text_repel` function from the `ggrepel` package.

#### Functional Enrichment Analysis with ShinyGO

To interpret the biological significance of the dysregulated genes, functional enrichment analysis is performed using the ShinyGO web tool [19]. The lists of significantly up-regulated and down-regulated genes (meeting the criteria of |*avg log*_2_*FC*| *>* 0.6 and *p val adj <* 0.05) extracted from the DE analysis are submitted to ShinyGO collectively. The analysis is run against the Gene Ontology (GO) Biological Process database. The resulting enrichment maps, which depict significantly over-represented biological pathways, are integrated into the study to provide a systems-level understanding of the functional alterations associated with PDAC in a cell-type-specific manner.

### 2.5 Cell Trajectory Analysis

To investigate the developmental trajectories and potential state transitions of immune cell populations in PDAC, we also perform single-cell trajectory analysis using the Monocle3 package. This approach models continuous biological processes, such as cell differentiation or activation, that underlie the discrete clusters identified in our data [20].

The analysis utilizes high-quality cells selected through confidence thresholds as described in section 2.3 (`predicted.celltype.l1.score` *>* 0.7 and `mapping.score` *>* 0.5), followed by dimensionality reduction with PCA, batch correction, and UMAP embedding. Trajectory graphs are constructed using the `learn_graph` function to infer potential developmental paths.

The resulting trajectories highlight distinct lineage relationships and transition patterns among major immune cell subtypes. In particular, the pseudotime ordering reveals progressive differentiation from naïve or precursor-like states toward activated and exhausted phenotypes, suggesting a dynamic reshaping of immune function within the PDAC microenvironment. Moreover, when cells are colored by fine-grained subtypes and faceted by condition (`“Cancer”` and `“Healthy”`), we observe condition-specific bifurcations, where certain immune lineages in tumor tissues appear to diverge into pro-tumorigenic or immunosuppressive branches.

This comparative trajectory analysis provides biological insight into how the immune landscape evolves during PDAC progression, illustrating that tumor-associated immune populations may follow altered developmental programs that potentially contribute to immune evasion and disease advancement.

## 3 Results

### 3.1 Data Integration and Cell Type Annotation

Figure 1 illustrates the cellular heterogeneity analysis of PBMC samples. Following quality control and filtering, Harmony integration successfully aligns cells from different samples, as evidenced by the intermixed distribution in the UMAP visualization colored by sample origin (Figure 1 A). We then annotate cell types based on the Azimuth reference-based classification. As detailed in section 2.3, only cells with a high-confidence Azimuth prediction are retained for visualization and downstream analysis. The resulting UMAP in Figure 1, B-D reveals the major immune populations according to the hierarchical classification system, including major cell types such as T cells, B cells, and monocytes, which all form distinct, well-separated clusters in the integrated space.

**Figure 1.**
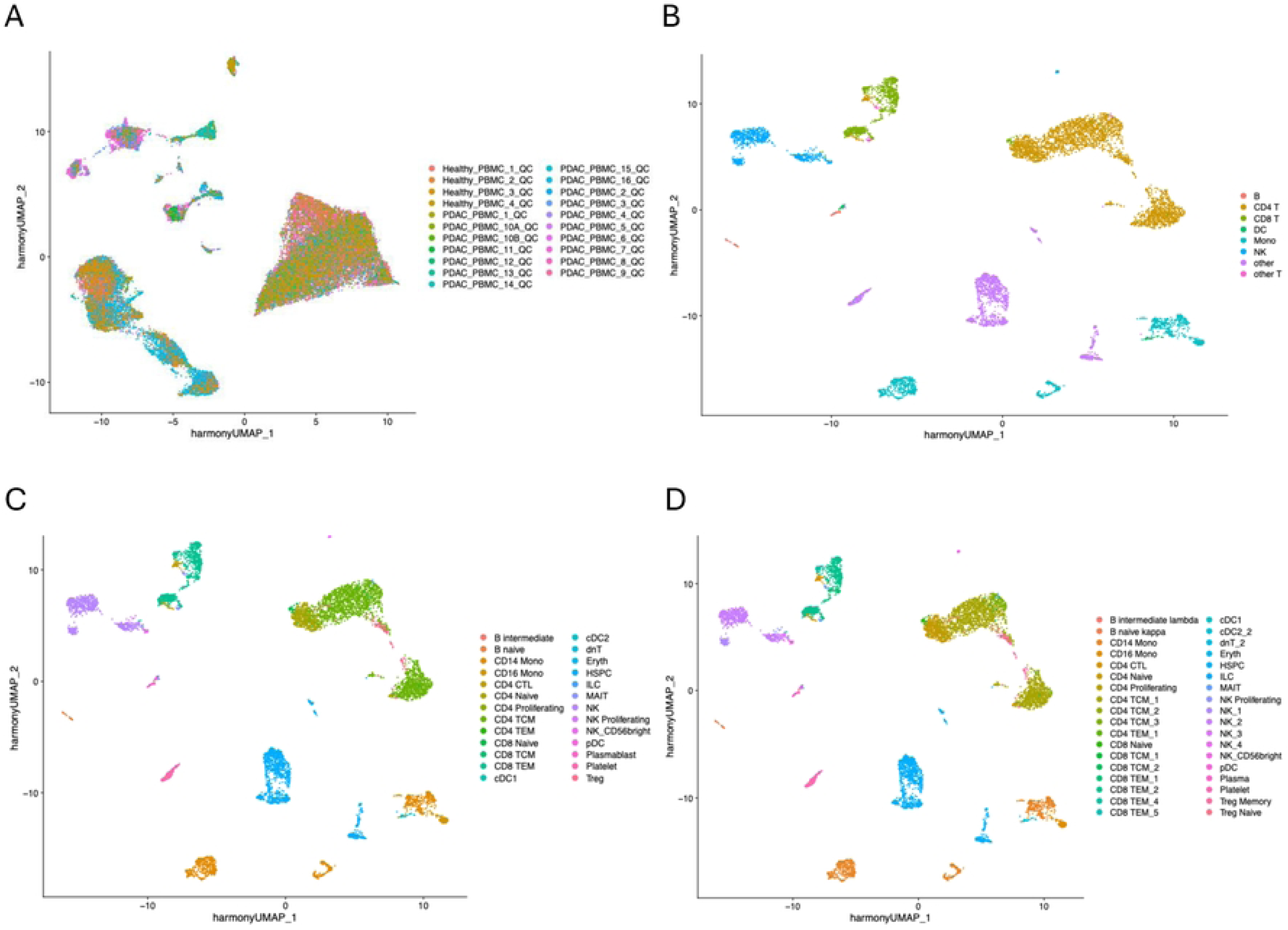

Figure 2 presents the parallel analysis of pancreatic tissue samples. Similarly, Harmony integration effectively corrects for batch effects, yielding a uniform distribution of cells from adjacent normal tissue and PDAC patients (Figure 2 A). Cell type annotation, performed using the same high-confidence Azimuth strategy described in section 2.3, identifies the major constituent cells of the pancreatic tissue microenvironment. Figure 2 B visualizes these annotated cell types, such as ductal cells, acinar cells, and various immune populations, which form clearly defined clusters in the integrated low-dimensional space.

**Figure 2.**
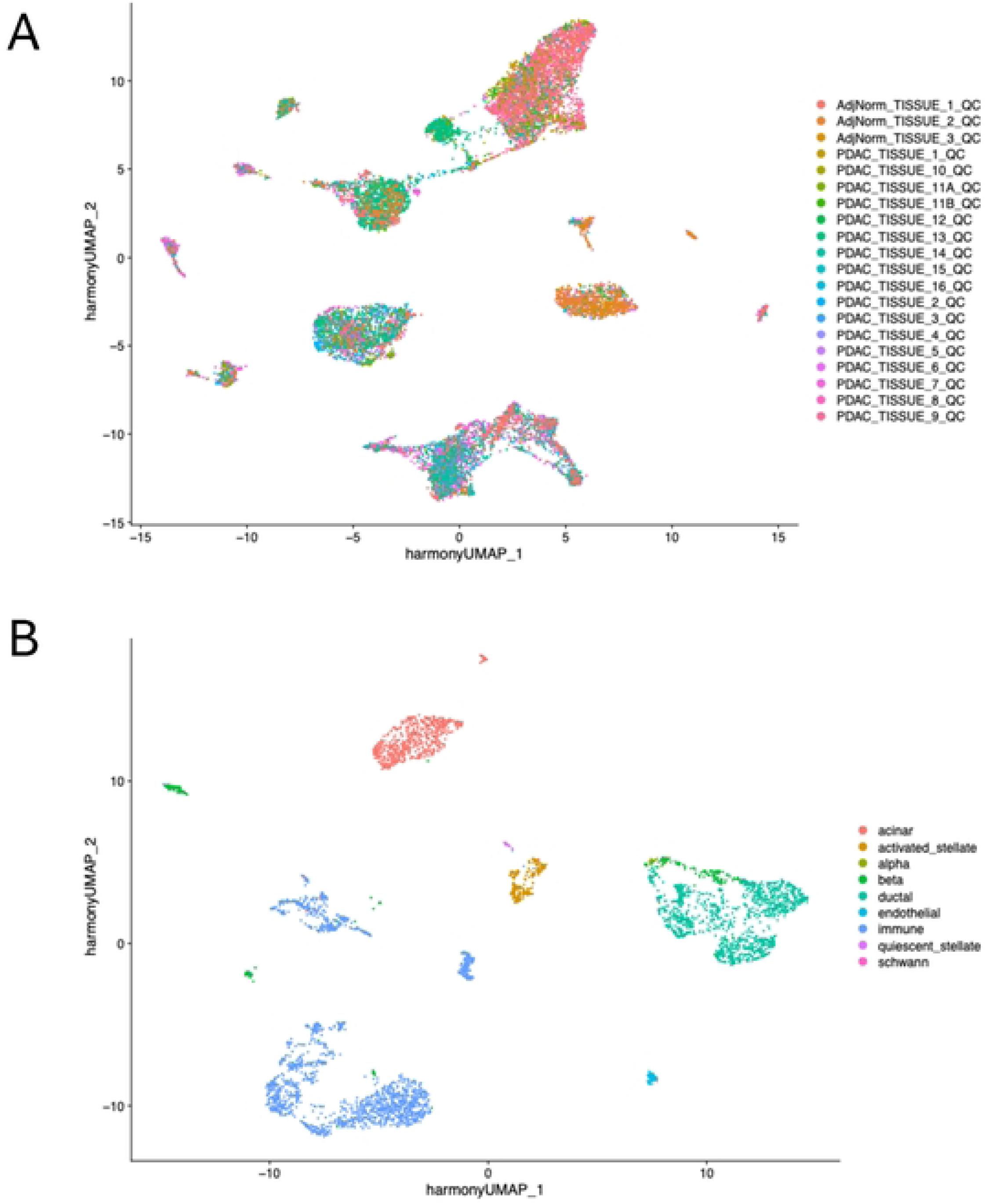

### 3.2 Differential Analysis

Following data integration and cell type annotation, we then perform comprehensive differential gene expression and pathway enrichment analyses to characterize transcriptional alterations in both pancreatic tissue and PBMCs from PDAC patients compared to healthy controls. Differential expression results are visualized using volcano plots that display log2 fold change on the x-axis versus −log10 adjusted p-value on the y-axis. Cyan points indicate significantly downregulated genes (log2FC *<* −0.6, adjusted p-value *<* 0.05), gray points represent non-significant genes, and red points show significantly upregulated genes (log2FC *>* 0.6, adjusted p-value *<* 0.05).

Pathway enrichment analysis uses ShinyGO with the complete human genome as background reference. The analysis visualizes the top enriched biological pathways where bar length indicates fold enrichment, which quantifies the overrepresentation of differentially expressed genes in a specific pathway compared to the expected proportion in the entire human genome. Color intensity represents −log10(FDR) values, where FDR denotes the false discovery rate adjusted p-value for pathway enrichment significance, with higher values indicating stronger statistical significance after multiple testing correction. Dot size shows the number of genes involved in each pathway.

#### 3.2.1 Pancreatic Tissue Analysis

Analysis of pancreatic tissue reveals cell-type specific alterations across epithelial and immune compartments. In acinar cells (Figure 3, A-B), we observe significant down-regulation of key function genes including *CELA2A* and *AMY2B*, alongside up-regulation of stress-response genes such as *RPL13* and *RPS3A*. Pathway analysis reveals enrichment in oxidative phosphorylation, metabolic pathways, and neurodegenerative disease-related pathways. Beta cells (Figure 3, C-D) show alterations in endocrine function genes including down-regulation of *CLPS* and *PRSS1*, and up-regulation of *CTRB2* and *CELA3B*, with corresponding pathway enrichment in pancreatic secretion, diabetes-related pathways, and various digestion and absorption pathways. Ductal cells (Figure 3, E-F) demonstrate significant changes in genes such as down-regulated *CLPS* and *PNLIP*, up-regulated *HSPB1* and *TMBIM1*, with pathway enrichment strongly focused on pancreatic secretion, fat and protein digestion and absorption pathways.

**Figure 3.**
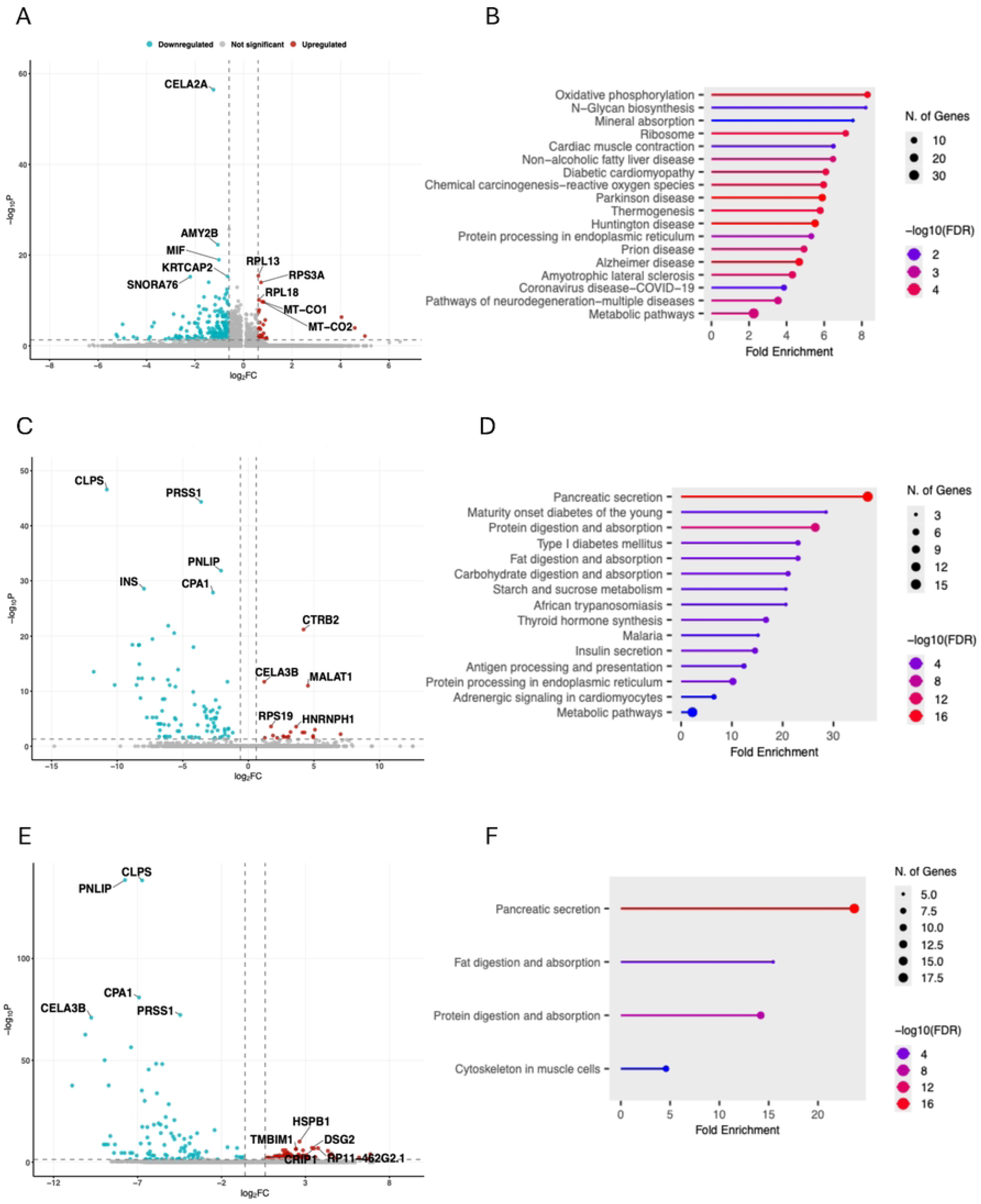

Within the pancreatic tissue immune compartment (Figure 4, A-B), immune cells show extensive transcriptional changes including up-regulation of *MIF* and *PPDPF*, and down-regulation of *CLPS* and *PRSS1*, with pathway enrichment in pancreatic secretion, digestive and metabolic pathways.

**Figure 4.**
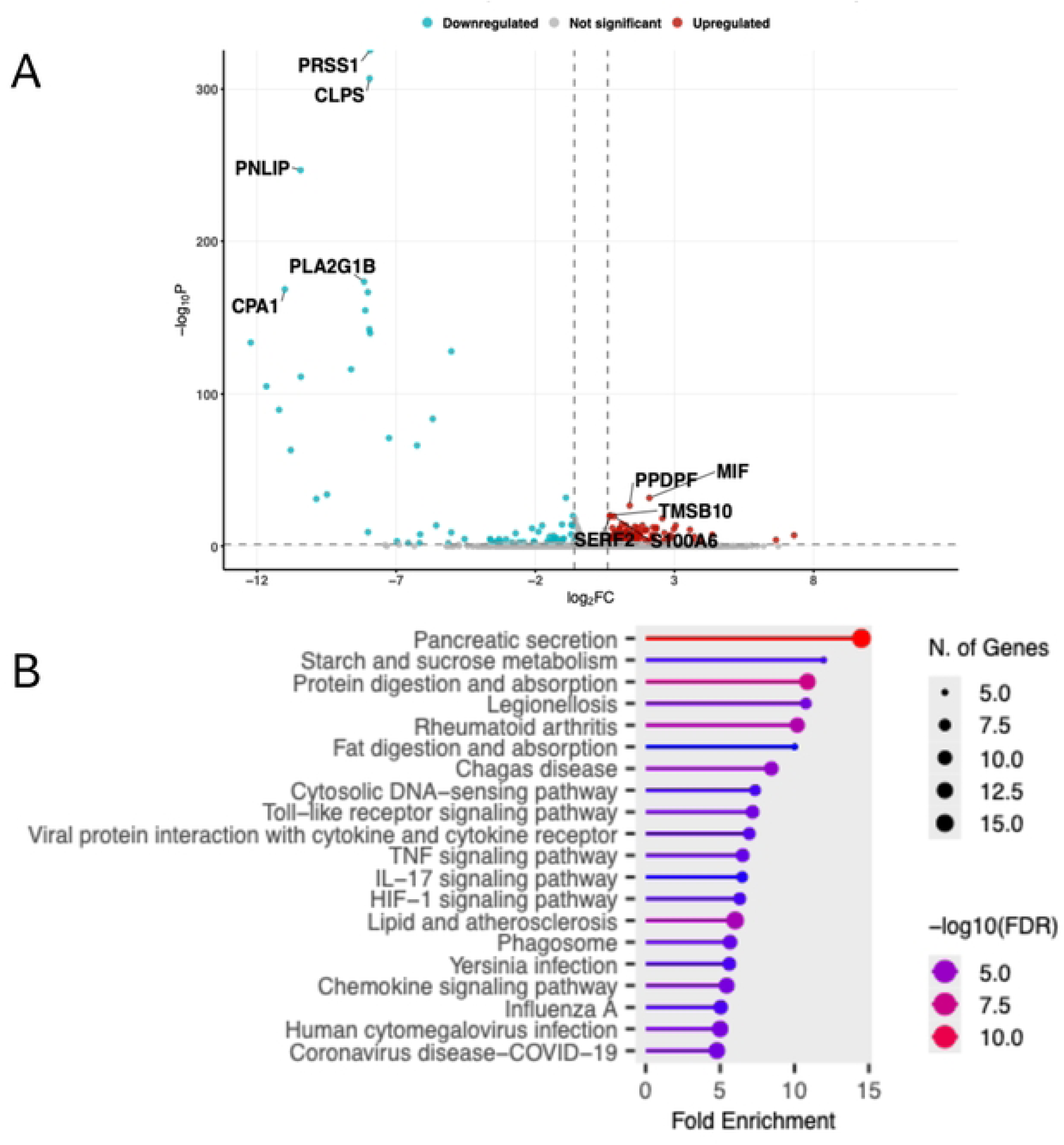

#### 3.2.2 PBMC Analysis

Analysis of peripheral blood mononuclear cells reveals systemic immune alterations across multiple cell lineages. Among cytotoxic populations, CD8 T cells (Figure 5, A-B) show activation of inflammatory genes including up-regulation of *ZFP36*, *CMC1* and *TSC22D3*, together with down-regulation of *MT-ND1*. Pathway analysis reveals enrichment in viral protein interaction with cytokine receptors, human cytomegalovirus infection, and cytokine-cytokine receptor interaction pathways. CD8 TEM cells (Figure 5, C-D) exhibit changes in up-regulated *ZFP36* and down-regulated *AKR1C3* expression, with pathway enrichment in malaria, leishmaniasis, and legionellosis pathways. NK cells (Figure 5, E-F) demonstrate alterations in down-regulated *RAB34*, *S100B* and *CX3CR1* genes, together with up-regulated *BTG1* genes, with extensive enrichment in diverse pathways including phototransduction, pertussis, renin secretion, and various metabolic and immune signaling pathways.

**Figure 5.**
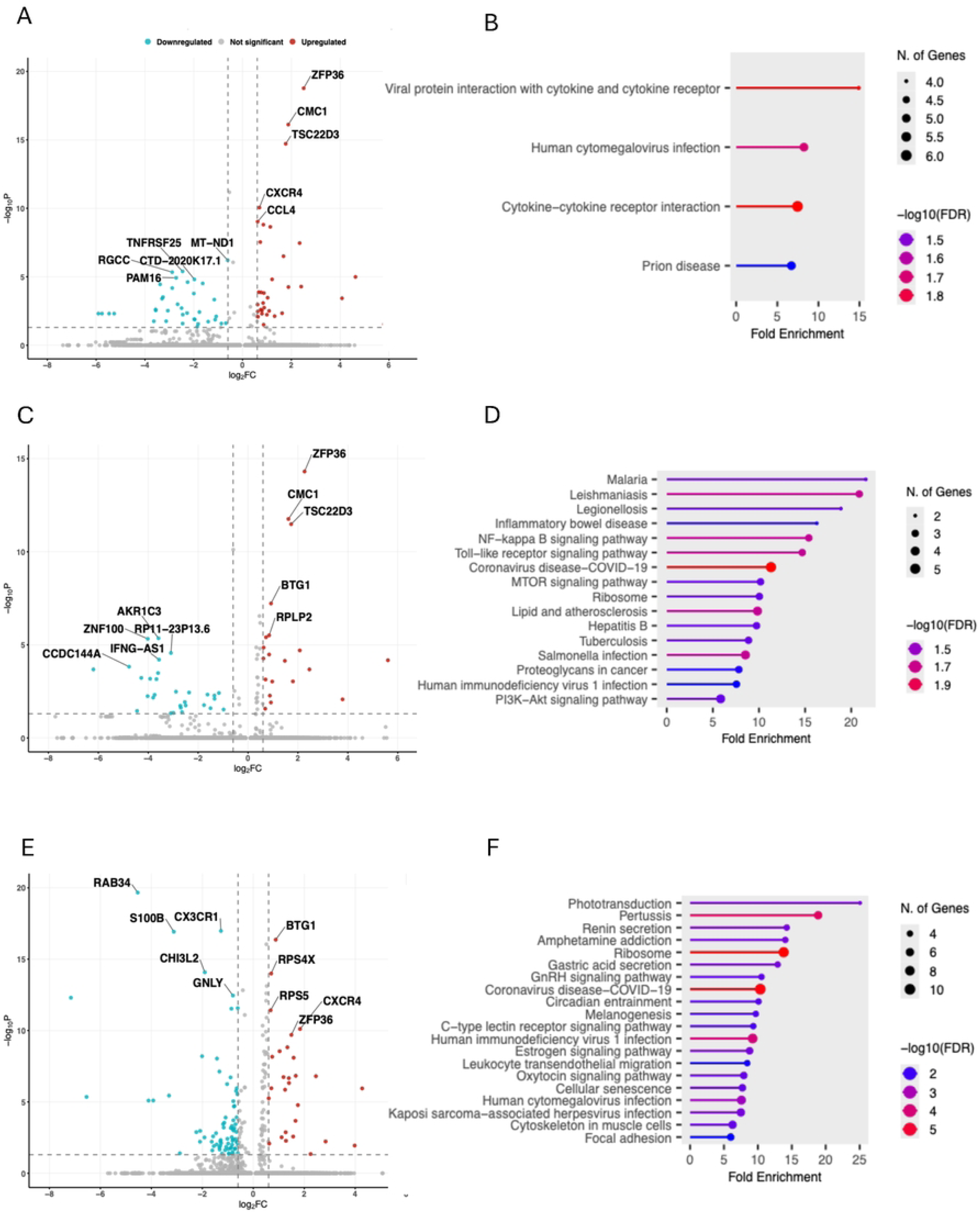

Among helper and memory T cell populations, CD4 T cells (Figure 6, A-B) display up-regulation of *RPL13A*, *NBEAL1* and down-regulation of *RPS4Y1*, accompanied by enrichment in IL-17 signaling pathway, Coronavirus disease-COVID-19, and Salmonella infection pathways. CD4 TCM cells (Figure 6, C-D) exhibit changes in up-regulated *RPL13A*, *RPL36A* and down-regulated *RPS4Y1* expression, with pathway enrichment in prostate cancer, toxoplasmosis, and infectious disease-related pathways.

**Figure 6.**
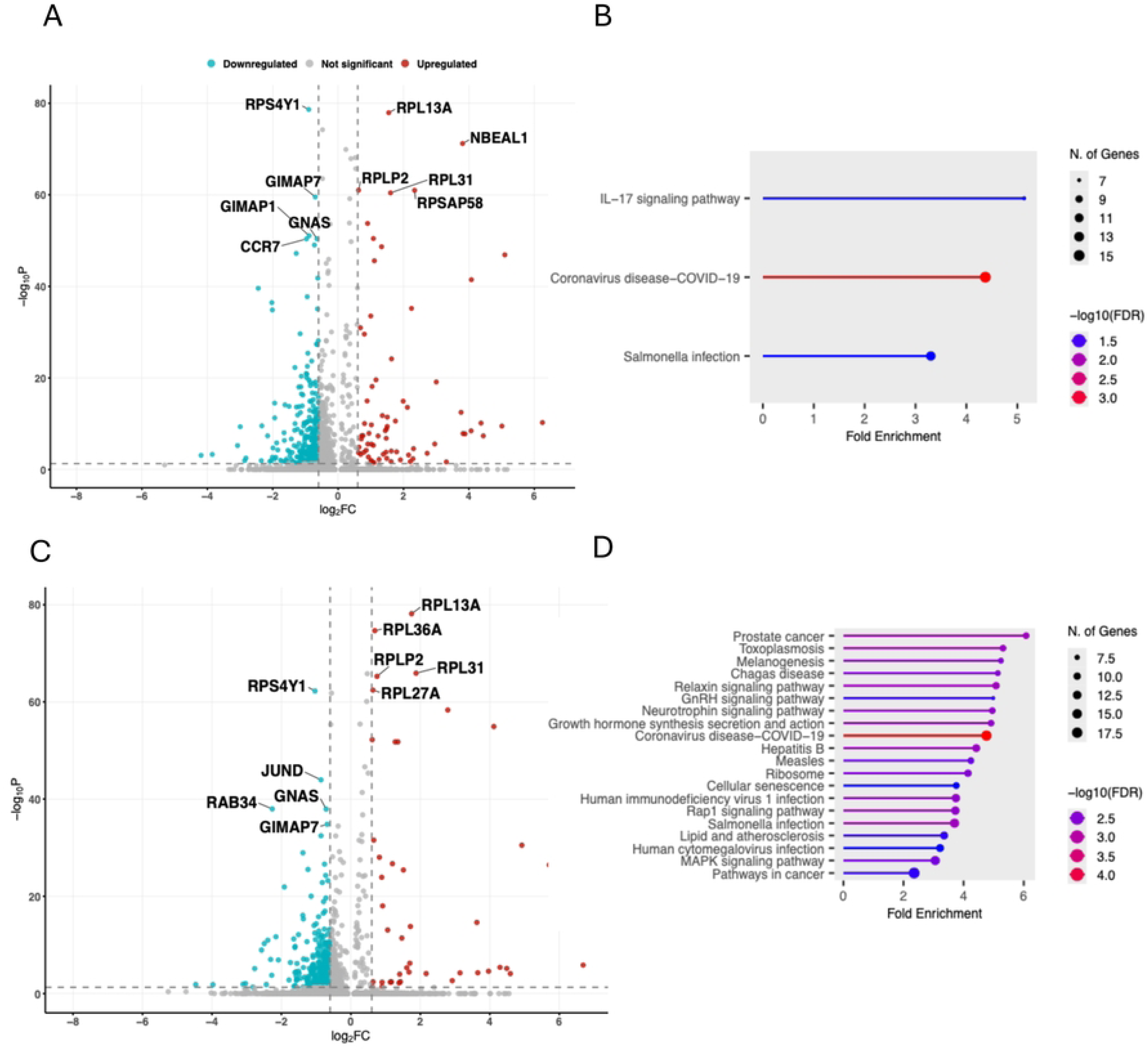

Myeloid and progenitor populations also show significant alterations. HSPCs (Figure 7, A-B) display up-regulation of *RPL32* and *RPS15*, and down-regulation of *SNCG*, *WFDC2* and *BPIFB1*, alongside enrichment in type I diabetes mellitus, allograft rejection, and infectious disease pathways. Monocytes (Figure 7, C-D) show down-regulation of *FLNB* and *HABP4* without significant upregulated genes, with enrichment in cytoskeleton and calcium signaling pathways.

**Figure 7.**
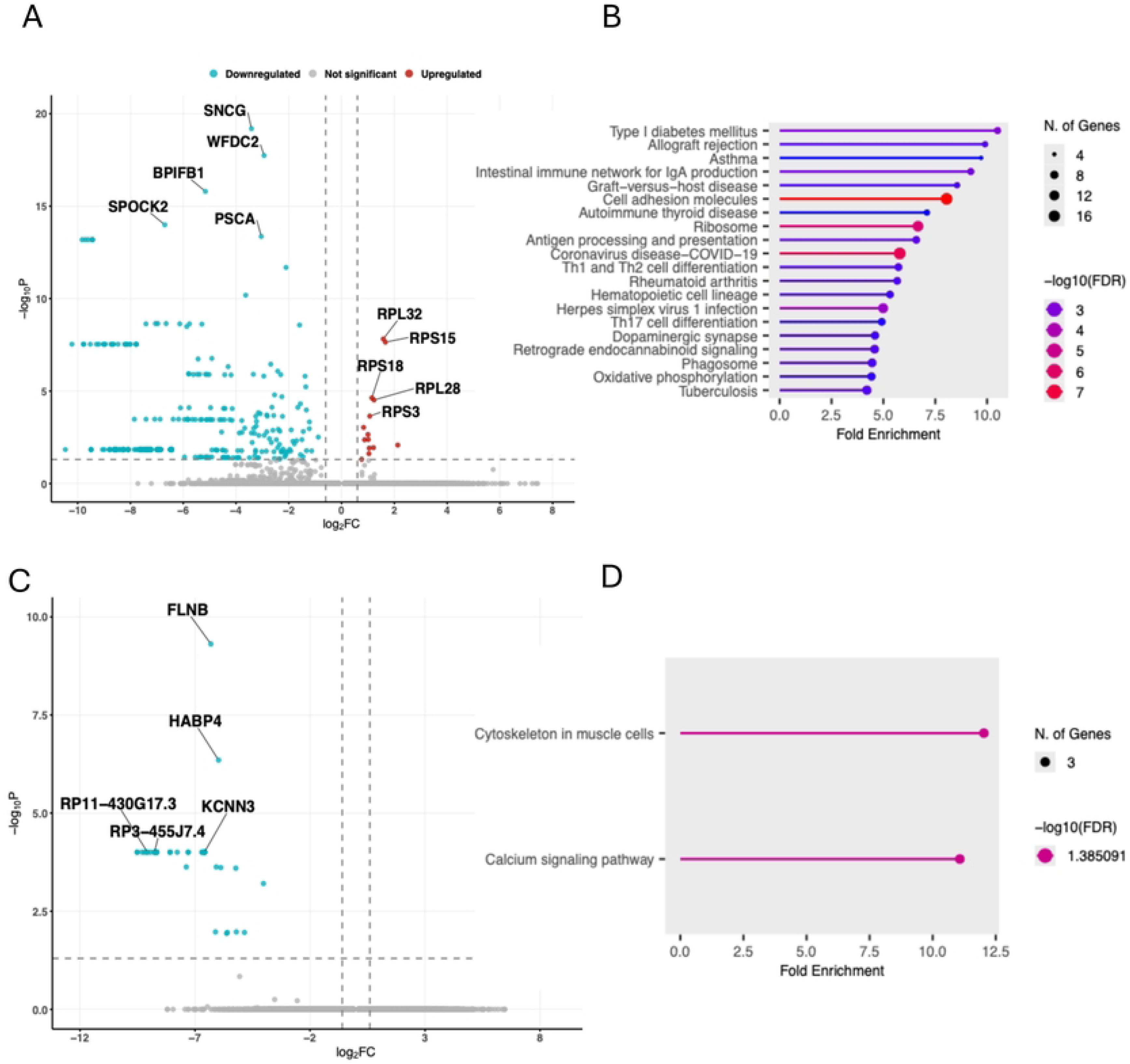

Humoral immune populations complete the PBMC characterization. B cells (Figure 8, C-D) show down-regulation of *RAB28*, *AKT1* and *EIF1AY* without significant up-regulated genes, accompanied by extensive enrichment in VEGF signaling, longevity regulating pathway, and numerous signaling cascades. Other unclassified cells (Figure 8, A-B) exhibit changes in *CD22*, *SNCG* and *RPS15* expression, with enrichment in allograft rejection, type I diabetes mellitus, and various immune and metabolic pathways.

**Figure 8.**
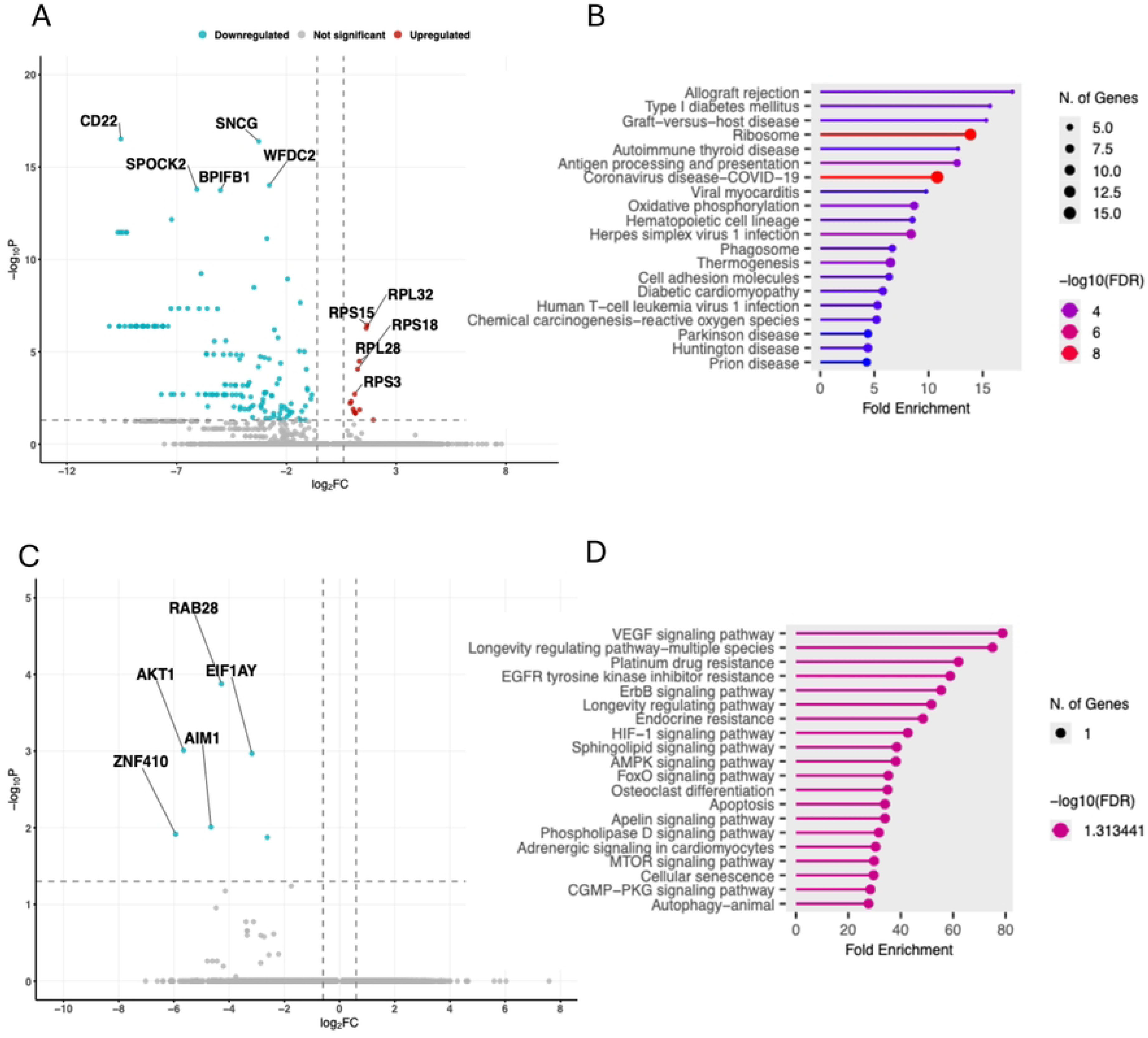

### 3.3 Comparative Trajectory Analysis of Immune Cell Subtypes

Cell trajectory analysis is further conducted on some cell types in PBMC samples to elucidate the dynamic differentiation pathways and fate decisions of immune cell sub-types under different pathological conditions. According to Figure 9 A, the faceted trajectory plot reveals distinct compositional differences in CD4 T cell subtypes between PDAC and healthy conditions within a conserved trajectory structure. In the PDAC microenvironment, CD4 T cells are predominantly comprised of CD4 TCM and CD4 CTL subsets, indicating a shift towards memory and cytotoxic phenotypes. In contrast, the healthy condition is characterized by a higher abundance of CD4 Naive cells, reflecting a more basal immunological state. This divergence in subtype distribution along shared developmental paths suggests that the tumor microenvironment promotes the differentiation of CD4 T cells towards specific effector and memory lineages, while preserving the overall trajectory architecture of CD4 T cell development.

**Figure 9.**
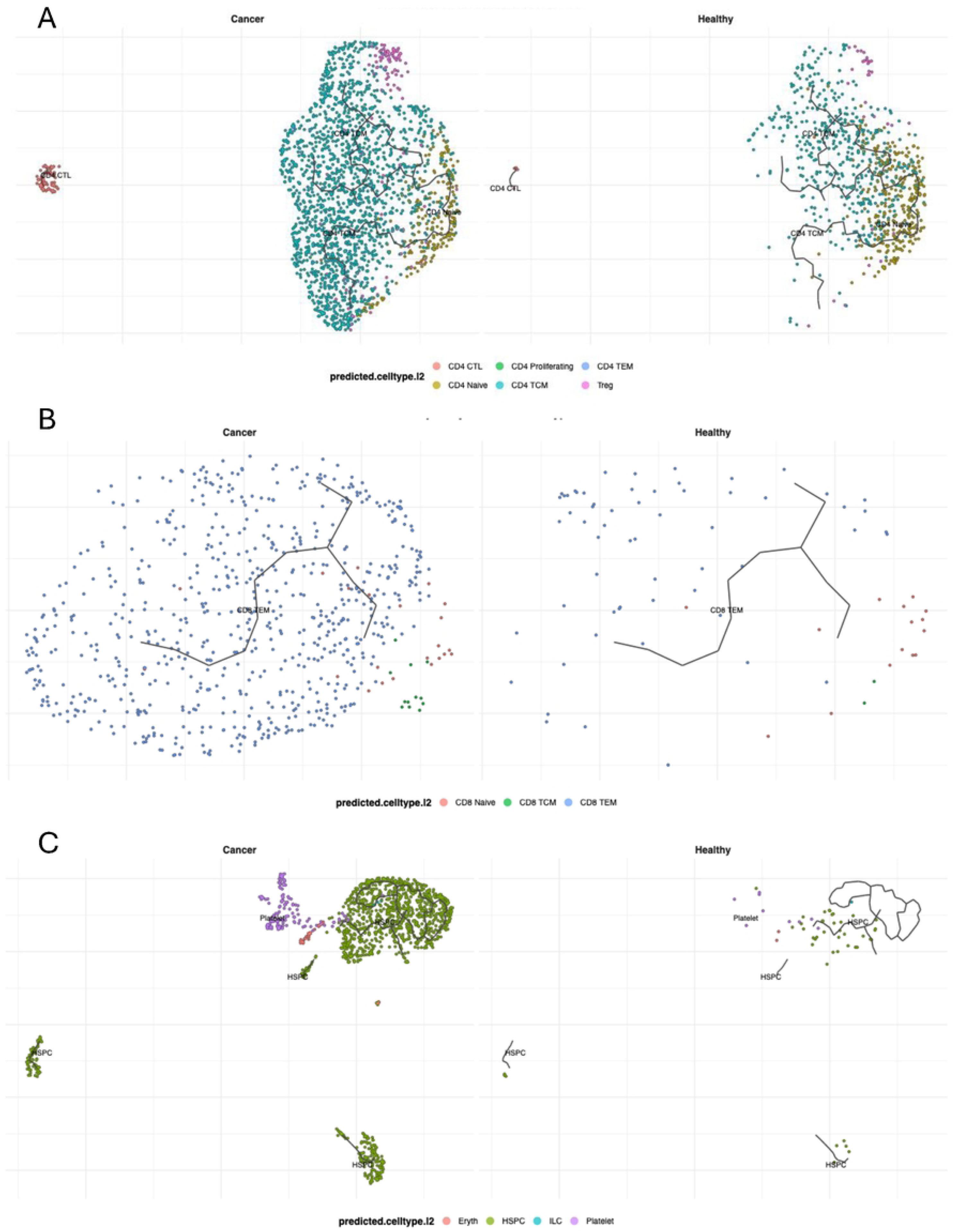

A similar trend was observed in CD8 T cells. Viewing from Figure 9 B, in the tumor microenvironment, CD8 T cells are predominantly composed of the CD8 TEM subset, accompanied by a smaller proportion of CD8 TCM cells, indicating an active differentiation toward effector memory phenotypes. Conversely, healthy conditions display a markedly different distribution, characterized by a high frequency of CD8 Naive cells and substantially fewer TEM cells. This distinct subtype distribution along conserved developmental trajectories suggests that the PDAC microenvironment drives CD8 T cell differentiation toward effector memory states, potentially reflecting chronic antigen exposure and T cell activation within the tumor immune landscape.

Beyond adaptive immune populations, we also observe notable changes in progenitor and innate immune cells. The faceted trajectory visualization reveals significant alterations in the composition of other immune cell populations between PDAC and healthy conditions as shown in Figure 9 C. In the tumor microenvironment, hematopoietic stem and progenitor cells (HSPC) and platelets are substantially expanded compared to healthy conditions. This prominent increase in HSPCs and platelets along the developmental trajectory suggests active hematopoiesis and megakaryocyte differentiation within the PDAC microenvironment. The enrichment of these progenitor and platelet populations may indicate a tumor-driven demand for increased immune cell production and potential roles in tumor-promoting processes such as coagulation, inflammation, and immune modulation.

Together, these findings across CD4 T, CD8 T, and other immune lineages consistently demonstrate that PDAC reshapes the cellular composition within conserved developmental trajectories, favoring effector, memory, and progenitor phenotypes that may collectively contribute to an immunosuppressive microenvironment.

## 4 Discussion

Our comprehensive single-cell analysis of pancreatic tissue and peripheral blood reveals that PDAC pathogenesis represents a multi-compartmental process involving epithelial dysfunction, local immune remodeling, and systemic immune exhaustion. Across the epithelial landscape, insulin- and enzyme-producing cells of the pancreas undergo fundamental identity alterations during tumorigenesis. We observe consistent shutdown of digestive enzyme genes across all three major epithelial cell types—acinar cells show decreased expression of *CELA2A* and *AMY2B*, while beta and ductal cells turn down *CLPS*, *PRSS1*, *PNLIP*, and *CPA1*. This coordinated loss of specialized function suggests an active “de-skilling” process that facilitates tumor initiation, consistent with single-cell evidence showing acinar-to-ductal reprogramming and metaplastic transition during early pancreatic neoplasia [21, 22].

Concurrently, epithelial cells undergo major metabolic reprogramming to support rapid proliferation. Acinar cells show markedly increased expression of mitochondrial genes *MT-CO1* and *MT-CO2*, indicating enhanced oxidative phosphorylation activity. Such mitochondrial hyperactivation has been linked to metabolic remodeling that sustains malignant transformation in PDAC [22]. In parallel, upregulation of ribosomal proteins (*RPL13*, *RPS3A*, *RPL18*, *RPS19*) across epithelial cell types suggests increased biosynthetic demand. Ductal cells also display elevated expression of survival and stress-response genes (*TMBIM1*, *HSPB1*), consistent with the adaptive epithelial states observed during oncogenic transformation [22].

In the immune compartment, the tumor microenvironment exhibits profound remodeling that paradoxically promotes cancer progression. We detect elevated levels of *MIF*, *TMSB10*, and *S100A6*, indicative of chronic inflammation and immune dysregulation. This aligns with reports that MIF-driven cytokine loops enhance immune evasion and fibrotic remodeling in PDAC [23]. Pathway analysis further reveals a paradoxical state in which digestive and tissue-remodeling pathways are highly upregulated while inflammatory cascades (TNF, IL-17, and Toll-like receptor signaling) remain persistently active. Sustained inflammation of this nature reprograms macrophages into tumor-supportive phenotypes and amplifies immune suppression [24]. In particular, *MIF* overexpression promotes angiogenesis and pro-survival signaling, fostering a microenvironment that favors malignant growth.

Beyond the local milieu, systemic immune dysfunction is evident in circulating cytotoxic compartments. CD8 T, CD8 TEM, and NK cells all display exhaustion signatures and impaired effector capacity. CD8 T cells activate viral-response pathways, reflecting antigenic distraction and energy misallocation, a phenomenon also observed in recent single-cell immune profiling of PDAC [25]. Effector molecules essential for tumor clearance, such as *CX3CR1* in NK cells and *TNFRSF25* in CD8 T cells, are downregulated, while the immunosuppressive regulator *TSC22D3* is upregulated, consistent with terminal T cell exhaustion [26]. Furthermore, mitochondrial imbalance (*CMC1* up, *MT-ND1* down) indicates compromised energy metabolism, reinforcing the presence of metabolic exhaustion [25].

Helper T cell subsets also show pronounced dysregulation. CD4 T cells exhibit overactivation of the IL-17 pathway and downregulation of the homing receptor *CCR7*, indicating an inflammatory yet spatially disoriented immune state. These findings echo spatial transcriptomic observations of disrupted T cell zonation and signaling networks within PDAC tissues [27]. Similarly, CD4 TCM cells display identity confusion and aberrant activation of cancer- and infection-related programs, coupled with reduced expression of transcriptional regulators (*JUND*, *GNAS*) and increased ribosomal activity—features characteristic of terminally differentiated and functionally exhausted memory T cells [27].

The myeloid lineage further reflects systemic derangement. Hematopoietic stem and progenitor cells (HSPCs) show aberrant activation of autoimmune programs while suppressing key defense genes (*SNCG*, *WFDC2*, *BPIFB1*), suggesting defective myelopoiesis driven by chronic inflammation. Similar stem-to-myeloid misdifferentiation and loss of phagocytic competency have been documented in recent single-cell PDAC atlases [28, 29]. Circulating monocytes also exhibit structural (*FLNB*), communication (*HABP4*), and activation (*KCNN3*) defects, consistent with the paralyzed, immunosuppressive phenotype of tumor-associated monocytes reported in PDAC [29].

Finally, B cell compartments exhibit a breakdown of humoral immunity. One subset shows autoimmune activation with loss of the inhibitory receptor *CD22*, consistent with reports that B cell dysregulation contributes to tumor-associated autoimmunity and chronic inflammation in pancreatic cancer [30]. Conventional B cells display widespread downregulation of genes involved in survival (*AKT1*), trafficking (*RAB28*), and translation (*EIF1AY*), suggesting apoptosis and loss of antigen-presenting capacity, consistent with recent observations of B cell exhaustion across multiple solid tumors including PDAC.

Taken together, our findings describe pancreatic cancer as a system-wide break-down in which epithelial cells lose identity and adopt metabolic resilience, while immune networks at both local and systemic levels collapse into dysfunction. Such cross-compartmental failure has been emphasized in recent integrative multi-omics reviews of PDAC, highlighting how epithelial reprogramming, fibroinflammatory signaling, and systemic immune exhaustion act as mutually reinforcing processes [24, 28].

## 5 Conclusion

This study highlights the power of integrative single-cell bioinformatics approaches in elucidating the complex immunometabolic landscape of pancreatic ductal adenocarcinoma (PDAC). By simultaneously capturing epithelial, immune, and systemic compartments, we establish a comprehensive framework that reveals how cellular cross-talk and lineage dynamics collectively drive tumor progression. These findings extend current understanding beyond traditional tumor-centric perspectives, offering new hypotheses on immune dysregulation and potential therapeutic intervention points. In particular, the identification of coordinated transcriptional programs across immune subpopulations provides a conceptual basis for developing targeted immunometabolic or precision therapies in PDAC. Although this work relies primarily on computational analyses without experimental validation, it nevertheless offers valuable insights and testable hypotheses that advance our understanding of PDAC biology and lay the groundwork for future translational research.

## Declarations

### Ethics approval and consent to participate

Not applicable

### Consent for publication

Not applicable

### Availability of data and material

All data and code generated or analyzed during this study are publicly available in the GitHub repository at https://github.com/Qinyu57/Bioinformatics-Analysis-of-Pancreatic-Cancer.

### Authors’ contributions

QX conceived and designed the study, performed all data analyses, prepared the figures, and wrote the manuscript. All aspects of the research, including data processing, interpretation, and manuscript revision, were completed independently by the author.

## Acknowledgements

The author would like to express sincere gratitude to Dr. Guangwei Zhang from the Keck School of Medicine, University of Southern California, for his valuable mentorship, insightful discussions, and continuous guidance throughout the course of this study. His expertise and encouragement greatly contributed to the completion of this work.

